# A *Plasmodium* lipase is critical for the disruption of the parasitophorous vacuole membrane and egress of hepatic merozoites

**DOI:** 10.1101/2025.06.02.657352

**Authors:** Raksha Devi, Pragya Mehra, Shabeer Ali, Satish Mishra

## Abstract

Malaria parasites develop within a specialized vacuole delimited by the parasitophorous vacuolar membrane (PVM). The exit of *Plasmodium* from host cells is a well-regulated process involving several molecules that disrupt the PVM to continue the life cycle. While several protein classes play a role in *Plasmodium* PVM rupture and merozoite egress, the first class of enzymes one would consider to act on membranes is lipases. However, the role of a lipase in PVM disruption remains unknown. Here, we characterized the role of UIS28 (upregulated in infective sporozoite gene 28) in *Plasmodium berghei.* Bioinformatic studies indicated that UIS28 is a class 3 lipase containing a fungal lipase-like domain. We found that UIS28 localizes to the PVM in infected hepatocytes. To understand the role of UIS28 in PVM disruption, we validated its lipase activity and found that it breaks down lipids into glycerol. Parasites lacking UIS28 develop normally in the blood and mosquito stages and mature fully into hepatic merozoites but show a defect in egress from host hepatocytes. We quantified the PVM rupture and reported that mutant parasites exhibit delayed egress due to impaired PVM disruption. Together, our results demonstrate that UIS28 is a lipase involved in the disruption of the PVM and the successful egress of hepatic merozoites.

**Author Summary:** An essential step in the life cycle of malaria parasites is their egress from hepatocytes, allowing them to transition from the liver stage to the blood stage. Liver stage parasites reside in a PV, which is surrounded by a PVM. Parasites must breakdown the PVM before they can infect other cells. This complex, multistep process is orchestrated by the parasite and involves several molecules to disrupt the PVM. However, little is known about these molecules. Here, we validated that UIS28 is indeed a lipase. We show that UIS28 is expressed in sporozoites and in the liver stage, and that it localizes to the PVM of parasites within hepatocytes. Disruption of UIS28 gene does not affect blood and mosquito stage development. However, mutant parasites show a defect in the disruption of the PVM and egress of hepatic merozoites. For the first time, we demonstrate the role of a lipase in PVM rupture during the release of liver-stage merozoites.

## Introduction

*The Plasmodium* life cycle alternates between a mammalian host and a mosquito vector. The infection begins when an infected female *Anopheles* mosquito inoculates sporozoites into the mammalian host while probing for a blood meal [1]. Sporozoites migrate from the inoculation site to the liver, where they invade the hepatocytes and form a parasitophorous vacuole (PV) by invaginating the hepatocyte plasma membrane [2]. Within this vacuole, the parasite undergoes repetitive rounds of closed mitosis, resulting in the formation of thousands of merozoites that infect erythrocytes after egress from the hepatocyte [3]. The egress of merozoites from the hepatocyte represents an essential step for the transition of parasites from the liver to the blood [2].

The PV is a specialized membrane-bound compartment, and the PV membrane (PVM) is a dynamic interface that physically separates the parasite from the host cell cytoplasm [2]. The PVM is initially formed from the host cell membrane, which is extensively remodeled by the parasite through the introduction of parasite proteins [2]. The composition of the PVM is not completely known; however, several PVM localized proteins play critical roles in parasite development [4], including protection from host immune response, nutrient uptake, and waste elimination [5–7]. EXP1 (Exported Protein 1) was the first protein identified as a resident of the PVM and was shown to interact with host apolipoprotein H (ApoH) and help access nutrients from the host [8]. The PVM localized pore-forming protein EXP2 is expressed in blood, salivary gland sporozoites, and liver stage parasites [9,10]. During the blood stage infection, the protein acts as a component of the translocon complex and plays a pivotal role in exporting the proteins into the erythrocyte cytosol [11,12]. EXP2 also acts as a nutrient channel and helps in the exchange of nutrients between the host and parasite [13]. Parasites lacking EXP2 have a reduced ability to invade hepatocytes [14]. UIS3 and UIS4 are among the early transcribed membrane protein (ETRAMP) family genes, which are expressed solely during the sporozoite and liver stages and are localized to the parasite’s PVM [15]. UIS3 and UIS4 KO sporozoites invade hepatocytes normally but fail to complete liver stage development [16,17]. It was shown that UIS3 sequesters host LC3 to avoid elimination by autophagy in hepatocytes [18]. UIS4 was found to interact with the host cell actin, and the parasite avoids elimination by suppressing filamentous actin formation [5].

After parasites mature in the liver, the PVM ruptures, and merozoites are released in the form of merosomes [19]. The merosomes are released after disrupting two membranes: the PVM and the host cell plasma membrane [20]. Several proteins have been implicated in the rupture of the PVM and the parasite membrane during egress. Proteases, lipases, and pore-forming proteins (PFPs) are actively involved in PVM rupture. Proteases can facilitate the egress by either degrading integral membrane proteins and cytoskeletal elements of the host cell or by activating downstream effector molecules [2]. Lipases can promote egress either by facilitating the digestion of membrane lipids, which leads to disruption of the membrane, or by acting as signaling molecules [21,22]. The pore-forming protein PLP1 was found to be involved in the rupture of vacuole and the host membrane in *Toxoplasma gondii*, whereas in *Plasmodium*, PLP2 and PLP1 are required for egress of gametes [23,24] and asexual blood stage parasites, respectively [25]. Serine-rich antigen (SERA) proteins, a group of cysteine/serine proteases, play a pivotal roles in parasite egress from erythrocytes and hepatocytes [26,27]. In RBCs, egress is initiated by the discharge of secretory components from the microneme and exoneme, which is caused by calcium-dependent protein kinase called CDPK5-dependent activation of cGMP-dependent protein kinase (PKG). The discharge of SUB1 (subtilisin-like parasite protease) into the PV lumen from the exoneme initiates a downstream cascade and activates the multiprotein complexes MSP1/6/7 and SERA6. SERA6 cleaves β-spectrin to destabilize the red blood cell cytoskeleton after PVM breakdown [28]. The role of PfSERA6/PbSERA3 in liver-stage egress, particularly whether it is required to destabilize the host cell cytoskeleton, remains to be determined.

LISP1 (liver-specific protein 1), a protein expressed during late liver stage development, was found to be involved in the disruption of the PVM and exit of the parasite from hepatocytes [29]. A member of the SERA family, SERA4, a cysteine protease, was reported to be critical for the breakdown of PVM and egress of merozoites from the hepatocyte [30]. The phospholipase, PbPla1 is critical for parasite egress from hepatocytes. Hepatic merozoites lacking PbPla1 showed reduced mobility and delayed parasite egress from hepatocytes [31]. Another PVM localized phospholipase, PbPL, was also found to be important for the disruption of the PVM and hepatic merozoite egress [20]. Despite an understanding of the role of proteases, kinases, and phospholipases in parasite egress, the precise role of a lipase in governing membrane disruption remains unknown.

In this study, we report that UIS28 is a lipase that is localized to the PVM of infected hepatocytes. By using a conventional knockout (KO), we showed that UIS28 is not required for parasite development in the blood or mosquito stages but is critical for the egress of hepatic merozoites. This is the first report of a protein with lipase activity implicated in PVM disruption during merozoite egress from hepatocytes.

## Results

### UIS28 is highly conserved within *Plasmodium* and contains a class-3 lipase domain

We started our study with the UIS28 amino acid sequence analysis. The BLAST search revealed the conservation within *Plasmodium* and considerable sequence similarity with *Hepatocystis sp* and *Eimeria brunetti* (Fig. S1A and Table S1). ScanProsite and InterPro prediction analysis revealed that UIS28 is a secreted mono/diacylglycerol lipase containing a fungal lipase-like domain or class-3 lipase domain. The structure prediction performed with i-Tasser has provided the five most reliable structures. Based on the C-Score (−1.79), RMSD (14.1±3.8Å), TM-score (0.48±0.15), and B-factor (near zero), model 1 was selected for the structural studies. The primary structure and 3D model of UIS28 show the orientation of the catalytic stretch with serine residue and the lipase domain (pink) located within the α/β-hydrolase fold. The α/β-hydrolase fold within the lipase domain spans from 672-923. There is a short catalytic stretch ranging from 802-814 (PYVIIFTGHSFGAS) in the specified region with a highly conserved catalytic Ser-811 (Fig. 1 and Fig. S1B). It is evident from the analysis performed using the InterProScan tool and TMHMM 2.0 that the protein might be of transmembrane nature with multiple short membrane-spanning domains associated with cytoplasmic and outer membrane regions. The basic architecture of the protein begins with a short membrane-bound stretch of 5-27 aa at the N-terminal, followed by a 28-802 aa long region, which protrudes out of the membrane (non-cytoplasmic). The 825-978 aa long region is invaginated into the cytoplasm Fig. S2.

**Fig. 1.**
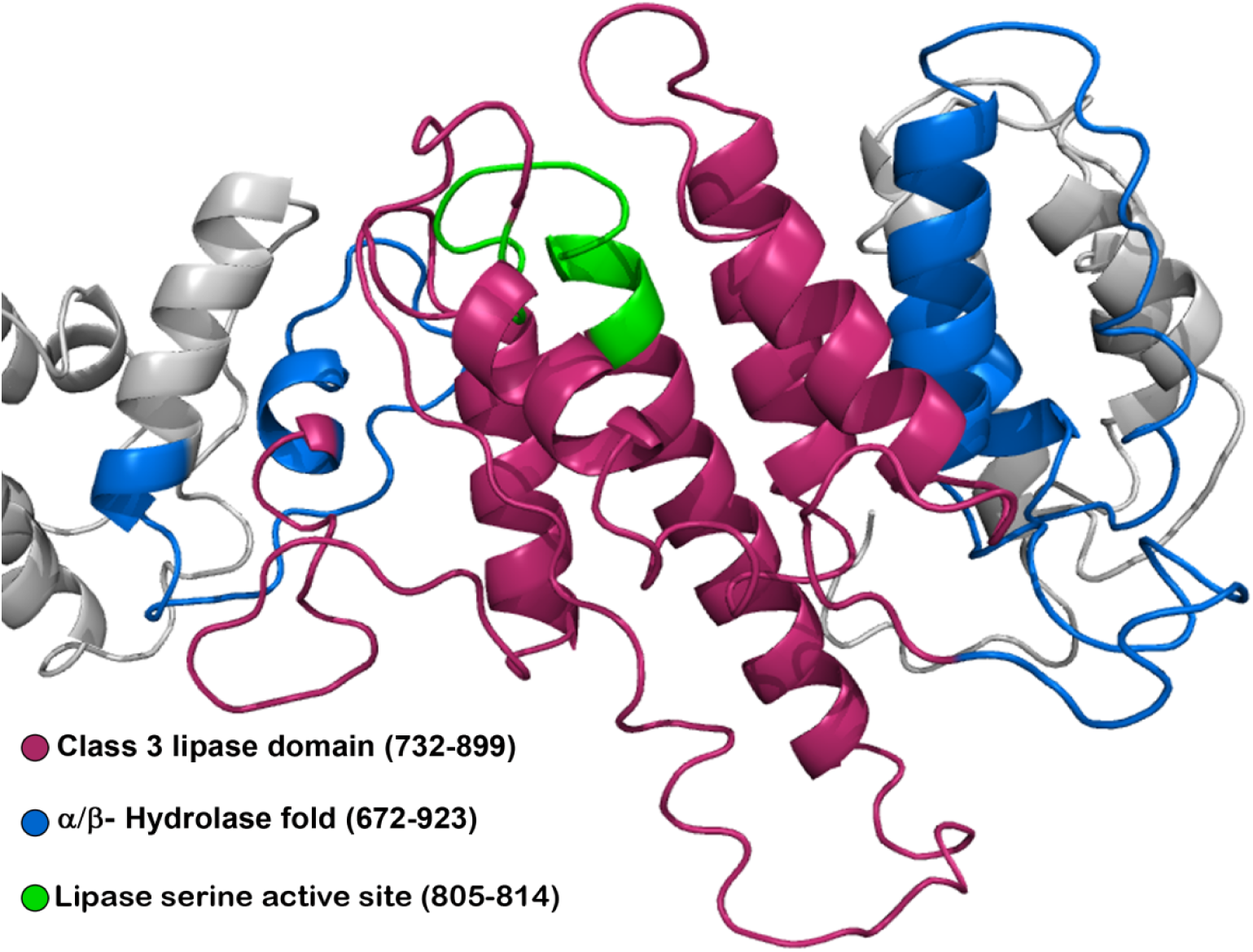
A portion of UIS28 showing the structural architecture of the class 3 lipase domain (warm pink) located within the a/b-hydrolase fold (blue colored region intervened by the pink region of the class 3 lipase domain). A small portion highlighted in green color is the lipase serine active site with the sequence PYVIIFTGHSFGAS.

### UIS28 is localized to the PVM of the liver stage

To investigate the expression and localization of UIS28, a transgenic parasite line expressing UIS28-3XHA-mCherry under the control of the endogenous promoter was generated (Fig. S3A). The correct integration of the targeting cassette was verified via diagnostic PCR (Fig. S3B). Next, we assessed the development of UIS28-3XHA-mCherry transgenic parasites (UIS28 Tr) throughout the parasite life cycle stages. We found that endogenous tagging of the gene did not affect UIS28 Tr parasite development in mosquitoes or mammalian hosts (Fig. S3C and D). For expression analysis, live parasites were examined under a fluorescence microscope. No mCherry signals were detected from asexual blood to mosquito midgut stages. We detected mCherry signals in salivary gland sporozoites and liver stages. The expression of the correctly sized UIS28-3XHA-mCherry fusion protein was confirmed in salivary gland sporozoites via western blot analysis. We detected an expected band size of ∼145 kDa in UIS28 Tr but not in the WT parasite lysate (Fig. 2A). Immunofluorescence assay (IFA) revealed that UIS28 is located throughout the sporozoites (Fig. 2B). The UIS28 expression pattern colocalized with UIS4 (PVM marker) in infected hepatocytes harvested at 48 and 65 h post-infection (hpi) (Fig. 2C). Overall, we conclude that UIS28 is a PVM protein.

**Fig. 2.**
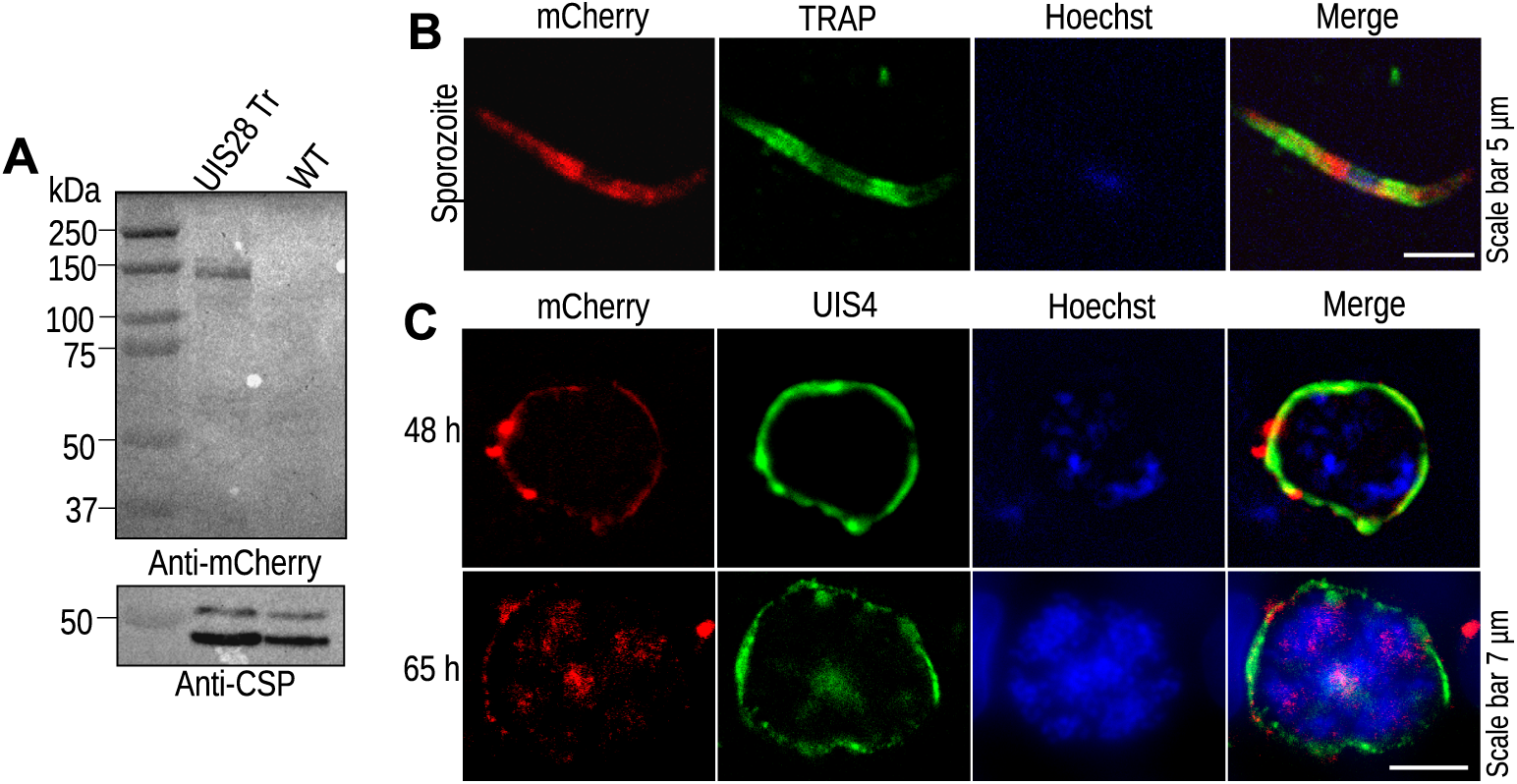
UIS28 localizes to the PVM. **(A)** Western blot analysis of the UIS28-3XHA-mCherry (UIS28 Tr) protein in salivary gland sporozoites via an anti-mCherry antibody. A band of ∼145 kDa was detected in UIS28 Tr parasites; no such band was detected in the WT parasite lysate. The membranes were stripped and reprobed with anti-CSP for detection of CSP as a loading control. **(B)** UIS28 Tr salivary gland sporozoites were immunostained with anti-mCherry and anti-TRAP antibodies. **(C)** Infected HepG2 cells were fixed at 48 and 65 h post infection (hpi) and immunostained with anti-mCherry and anti-UIS4 antibodies.

### UIS28 is a lipase

Bioinformatic analysis and IFA revealed that UIS28 is a class 3 lipase and is localized to the PVM. Next, we asked whether UIS28 indeed has lipase activity and is involved in PVM disruption. To test this hypothesis, we evaluated the lipase activity of *P. berghei* UIS28. We attempted to express the full-length protein in bacteria, but failed. Next, we successfully expressed the catalytic domain of UIS28 as a GST fusion protein (Fig. 3A). The expression of fusion protein was confirmed by western blot analysis via an anti-GST antibody, which recognized a band corresponding to the expected size of the fusion protein (Fig. 3B). Lipase activity in bacteria expressing GST-UIS28 was determined via an in vitro lipase assay kit. The UIS28 lipase activity was 1.9 milliunits/ml (Fig. 3C). No activity was observed with the negative control expressing GST alone. Taken together, these data strongly suggest that UIS28 is indeed a lipase.

**Fig. 3.**
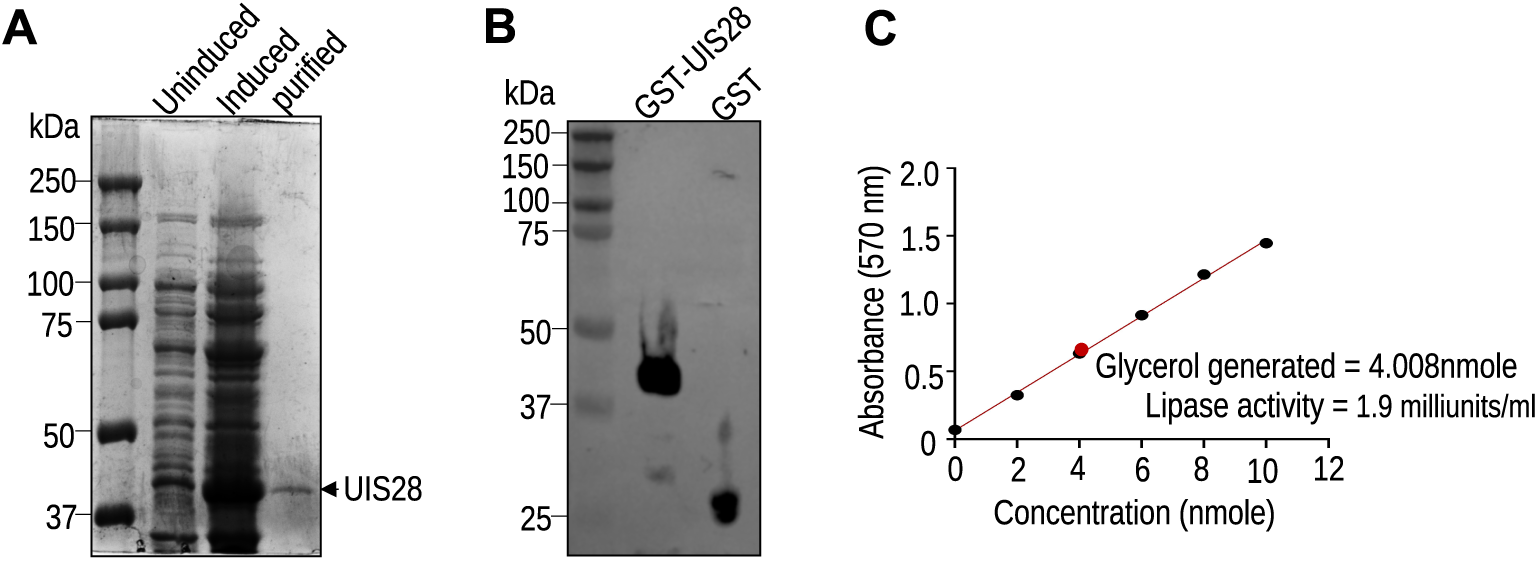
Expression and purification of UIS28 recombinant protein and lipase assay. **(A)** Coomassie brilliant blue-stained SDS-PAGE gel showing uninduced, induced, and purified GST-UIS28 protein. **(B)** Western blot analysis of recombinant GST-UIS28 using anti-GST antibody. A band of ∼44 kDa corresponding to the size of the GST-fusion protein was detected in the recombinant GST-UIS28 bacterial lysate compared with the 25 kDa GST control. **(C)** Determination of UIS28 lipase activity using GST-UIS28 fusion protein-expressing bacteria. The UIS28 protein generated 4.008 nmol of glycerol after 45 minutes of incubation with the lipase substrate at 37^°^C. The UIS28 lipase activity was 1.9 milliunits/ml.

### UIS28 is dispensable in *P. berghei* blood and mosquito stages

To study the role of UIS28 in *Plasmodium*, the gene was disrupted via a double crossover (DCO) homologous recombination strategy (Fig. S4A). After DCO, the UIS28 ORF was replaced with GFP and an hDHFR:yFCU expression cassette. Observation of GFP expressing blood stage parasites after drug selection confirmed successful transfection (Fig. S4B). Clonal lines (UIS28 KO c1 and c2) were obtained by limiting dilution of the parasites. Correct site-specific integration of the targeting cassette and the absence of the UIS28 ORF in the clonal lines were confirmed by diagnostic PCR (Fig. S4C). A reverse strategy was also employed to restore the gene function via complementation (UIS28^Comp^) to confirm the specificity of the KO phenotype (Fig. S4D). The restoration of the UIS28 locus was confirmed by diagnostic PCR (Fig. S4E). To investigate the effect of the loss of UIS28 on blood stage parasite growth, Swiss albino mice were intravenously injected with equal numbers of iRBCs. We detected comparable blood stage propagation of WT, UIS28 KO and UIS28^comp^ parasites (Fig. S4F). Next, the effect of UIS28 deletion on parasite development in the mosquito vector was analyzed. We infected female *Anopheles stephensi* mosquitoes by allowing them to feed on mice infected with WT or KO parasites. The development of the parasites was subsequently evaluated by quantifying ookinete formation, midgut oocyst development, and the generation of both midgut and salivary gland sporozoites. All the developmental stages of UIS28 KO parasites in mosquitoes were comparable to those of the control (Fig. S5A-H). Collectively, these results indicate that UIS28 is not required for the asexual blood stage propagation or mosquito stage development.

### UIS28 is critical for the parasite transition from liver to blood stage infection

To evaluate the in vivo infectivity of UIS28 KO sporozoites, C57BL/6 mice were intravenously injected with 5,000 sporozoites along with WT and complement parasites, and blood stage infection was observed via Giemsa-stained blood smear. All the mice infected with control parasites became patent on day 3 post-infection (pi). Compared with control, mice infected with UIS28 KO parasites consistently presented a 2-day delay in the appearance of blood-stage infections (Table 1). To identify the stage-specific defects, livers from another group of mice injected with sporozoites as described above were harvested at 40 and 72 h pi. To quantify the parasite load in the liver, RNA was isolated, cDNA was synthesized, and Pb18s rRNA was quantified via real-time PCR. At 40 hpi, there was no significant difference in parasite load between the WT and KO parasites (Fig. 4), indicating that the KO sporozoites are capable of infecting hepatocytes normally. By 60 hpi, liver parasites egress from the liver and initiate blood-stage infection. The parasite burden in livers infected with WT parasites was significantly lower than that in those infected with KO parasites at 72 hpi (Fig. 4). The greater parasite burden in the KO at 72 hpi indicates a defect in the egress of liver stage parasites.

**Fig. 4.**
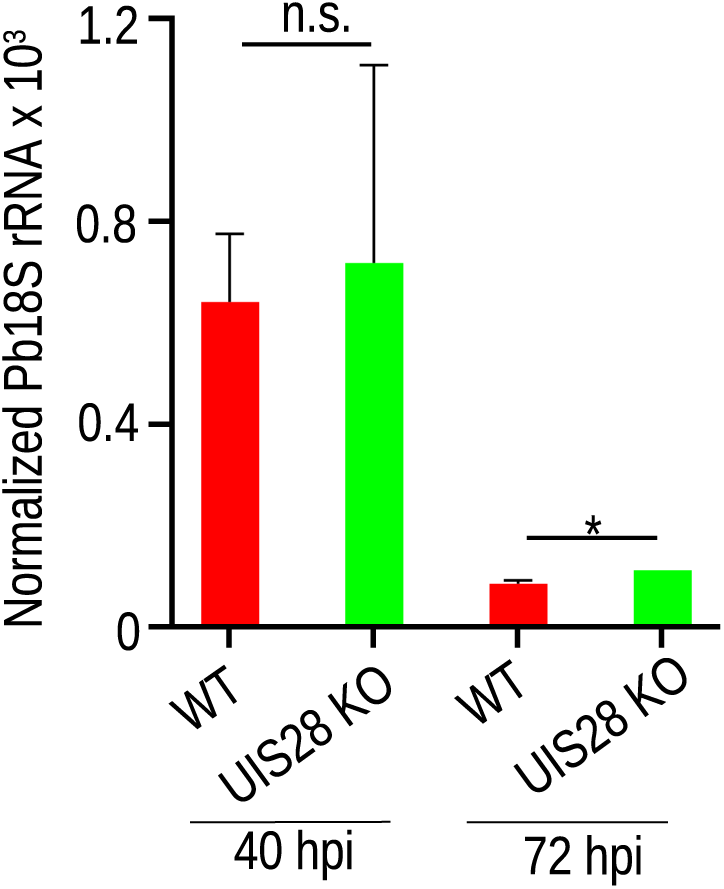
UIS28 KO parasites infect the liver but fail to egress efficiently. C57BL/6 mice were infected with sporozoites at 40 and 72 hpi. The mouse group that was used for harvesting the liver at 72 hpi was treated with chloroquine and artesunate, following sporozoite injection to kill the blood stage parasites. Livers were harvested, RNA was isolated, cDNA was synthesized, and the parasite load in the liver was quantified via real-time PCR via amplification of Pb18s rRNA. The transcripts were normalized to the levels of mouse GAPDH. No difference in transcript levels was observed at 40 hpi (P=0.8694). However, a significantly greater level of transcripts was detected in UIS28 KO parasite-infected livers at 72 hpi than in WT control parasites (*P=0.0161). The data are presented as the mean ± SEM from two (40 h) and three (72 h) independent experiments. N = 5 mice/group. The data were analyzed via Student’s t-test.

**Table 1.** In vivo infectivity of UIS28 KO sporozoites in C57BL/6 mice.

| Exp. | Parasite | Number of sporozoites inoculated | Mice positive/Mice inoculated | Prepatent period (days) |
| --- | --- | --- | --- | --- |
| **1** | WT GFP | 5,000 | 5/5 | 3.2 |
|  | PbUIS28 KO c1 | 5,000 | 5/5 | 5 |
|  | PbUIS28 KO c2 | 5,000 | 5/5 | 5.4 |
|  | PbUIS28 Comp | 5,000 | 5/5 | 3 |
| **2** | WT GFP | 5,000 | 5/5 | 3 |
|  | PbUIS28 KO c1 | 5,000 | 5/5 | 5.2 |
|  | PbUIS28 KO c2 | 5,000 | 5/5 | 5 |
|  | PbUIS28 Comp | 5,000 | 5/5 | 3 |

### UIS28 KO parasites mature into hepatic merozoites but exhibit a defect in egress

To further elucidate the role of UIS28 in liver stage development, the intrahepatic progression of KO parasites was examined in detail under in vitro conditions. Initially, the ability of UIS28 KO sporozoites to invade hepatocytes was assessed via an inside/outside assay. We found no significant difference in the invasion efficiency between WT and KO sporozoites (Fig. S6), further supporting our in vivo observations. Subsequently, HepG2 cells infected with UIS28 KO sporozoites presented a similar growth pattern and number of exoerythrocytic forms (EEFs) similar to those of WT GFP at 45 and 65 hpi (Fig. 5A-C). The EEF area of liver stage parasites was also determined at 45 and 65 hpi. No significant difference in size was observed between WT GFP, UIS28 KO, and UIS28 comp parasites (Fig. 5D). Furthermore, the development of KO parasites during late liver stages was investigated by immunostaining EEFs at 65 hpi with an anti-MSP1 antibody, which revealed that the cytomere and merozoite stages in UIS28 KO parasites were morphologically indistinguishable from those in control EEFs (Fig. 6A). The numbers of cytomere and merozoite stages were also counted, and there were no differences between the control and KO parasites at 55 hpi (Fig. 6B). However, number of merozoites were significantly decreased in WT GFP compared to UIS28 KO parasites at 72 hpi (Fig. 6B). The pattern of nuclear division in the KO parasites was comparable to that observed in the control parasites (Fig. 6C). The PVM surrounding the EEF in the liver must rupture to release merozoites into the bloodstream in the form of merosomes (detached cells in vitro). Compared with that in WT parasites, quantitative analysis of detached cells revealed a significant reduction in the number of detached cells produced in UIS28 KO parasites. Even longer incubation of the culture did not increase the number of detached cells in UIS28 KO parasites (Fig. 6D). The infectivity of detached cells was assessed in Swiss albino mice via intravenous injection of equal numbers of cells. The detached cells derived from WT and KO parasites exhibited similar infectivity (Table 2). Additionally, detached cells were immunostained with an anti-UIS4 antibody to evaluate PVM integrity. No intact PVMs were observed in either WT or UIS28 KO detached cells (Fig. 6E). Overall, these data suggest that UIS28 is required for hepatic merozoite egress.

**Fig. 5.**
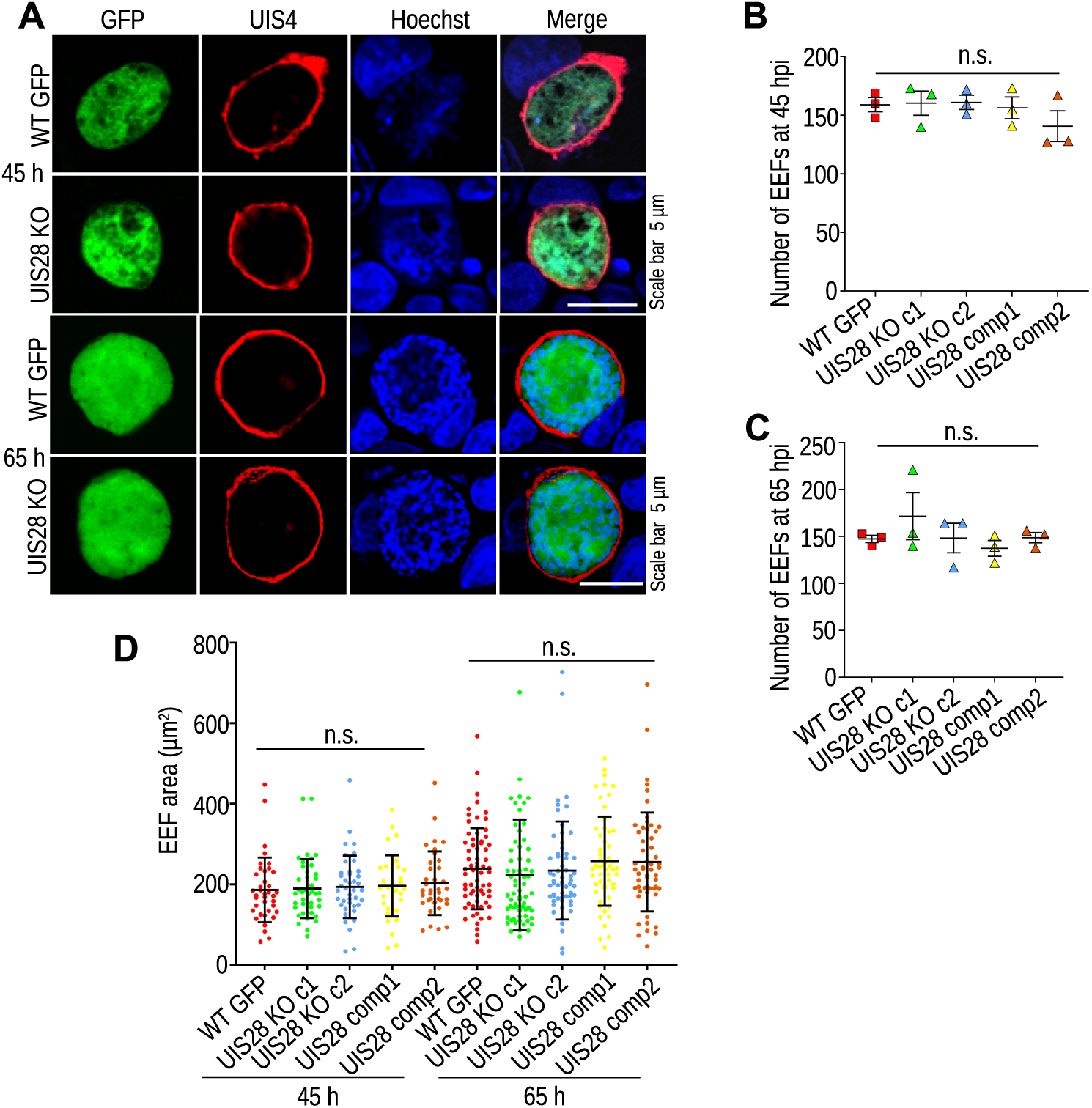
UIS28 KO parasites exhibit normal liver-stage development. **(A)** WT GFP and UIS28 KO infected HepG2 cultures were fixed at 45 and 65 hpi, and immunostained with an anti-UIS4 antibody. Nuclei were stained with Hoechst. **(B and C)** The number of EEFs produced in WT GFP and UIS28 KO cultures was counted at the indicated time points under a fluorescence microscope. No significant difference in EEF number was observed between WT and KO parasites, at 45 (P=0.5429) and 65 hpi (P=0.5460). The data are presented as mean ± SEM from three independent experiments performed in duplicate. **(D)** WT GFP and UIS28 KO EEF images were acquired via fluorescence microscopy, and the area was measured via density slicing using ImageJ. No significant difference in the growth of KO EEFs compared with WT EEFs was detected at 45 (P=0.2852) and 65 hpi (P=0.3961). A total of 38, 41, 41, 35, and 36 (45 hpi) and 67, 67, 60, 55, and 55 (65 hpi) EEFs of WT GFP, UIS28 KO c1, UIS28 KO c2, UIS28 comp1, and UIS28 comp2 parasites, respectively, were analyzed. The data are presented as mean ± SEM from three independent experiments. One-way ANOVA was used to analyze the statistical significance.

**Fig. 6.**
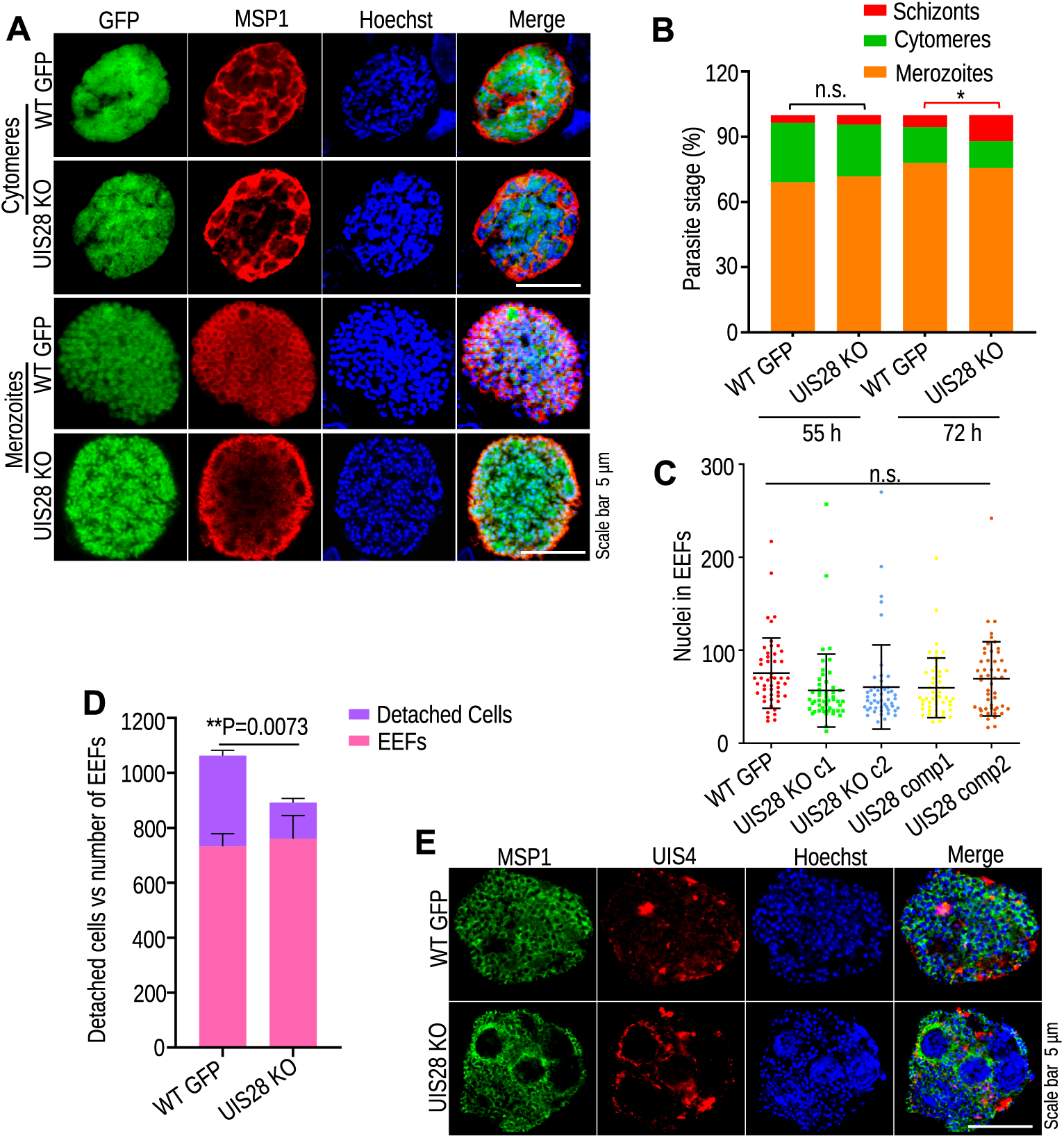
UIS28 is critical for the parasite egress from the hepatocytes. **(A)** WT GFP and UIS28 KO infected HepG2 cultures fixed at 65 hpi were immunostained with an anti-MSP1 antibody. The formation of the cytomere and merozoite stages was indistinguishable between WT GFP and KO parasites. **(B)** Schizont (P=0.5679), Cytomere (P=0.4844) and merozoite (P=0.0996) stage numbers were comparable in WT GFP and UIS28 KO parasites at 55 hpi. Schizonts (P=0.6564) and Cytomere (P=0.4149) were comparable in WT GFP and UIS28 KO parasites at 72 hpi, whereas merozoite numbers were significantly decreased in WT GFP parasites compared to UIS28 KO (*P=0.0124). **(C)** The number of nuclei in infected WT and KO EEFs was not significantly different (P=0.0962, one-way ANOVA). We analyzed 50 EEFs each for WT GFP, UIS28 KO c1, UIS28 KO c2, UIS28 comp1, and UIS28 comp2 parasites. The data were obtained from three independent experiments and are expressed as the mean ± SEM. **(D)** Detached cells were collected from the culture supernatant, counted via a hemocytometer, and normalized to the EEF count. The number of detached cells produced by UIS28 KO parasites was significantly lower than that produced by WT GFP parasites (P=0.0216, Student’s t test). The data were pooled from three independent experiments and are shown as the mean ± SEM. **(E)** Detached cells present in culture supernatants were immunostained with anti-MSP1 and anti-UIS4 antibodies. Nuclei were stained with Hoechst. The MSP1 and UIS4 staining patterns were similar in the WT and KO parasites.

**Table 2.**
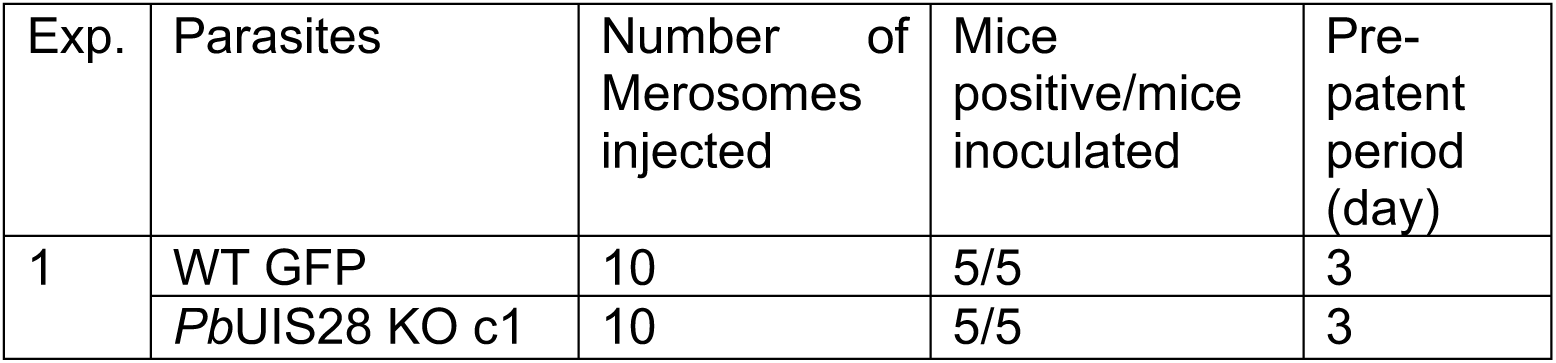
*PbUIS28* KO merosomes exhibit normal in vivo infectivity.

### UIS28 KO parasites exhibit a defect in PVM disruption

After the maturation of hepatic merozoites, the PVM ruptures, and liver-stage parasites are released into the host cell cytoplasm. A defect in PVM disruption leads to a reduced production of detached cells. We quantified PVM disruption by immunostaining UIS28 KO and WT EEFs with the PVM markers anti-UIS4 and anti-EXP1 antibodies at 65 hpi. We visually inspected images of EEFs and counted the number of EEFs with intact or disrupted PVMs. The percentage of EEFs with ruptured PVMs was significantly lower in the KO EEFs than in the WT EEFs (Fig. 7A-C). The breakdown of the PVM is accompanied by the collapse of the host cell’s actin cytoskeleton into the parasite cytoplasm. Next, we immunostained EEFs with an anti-actin antibody to analyze the host actin status in infected cells. Compared with those in control parasites, fewer KO EEFs were observed with actin present in the parasite cytoplasm, which is consistent with the reduced disruption of the PVM (Fig. 7D and E). These results establish a key role for UIS28 in PVM disruption.

**Fig. 7.**
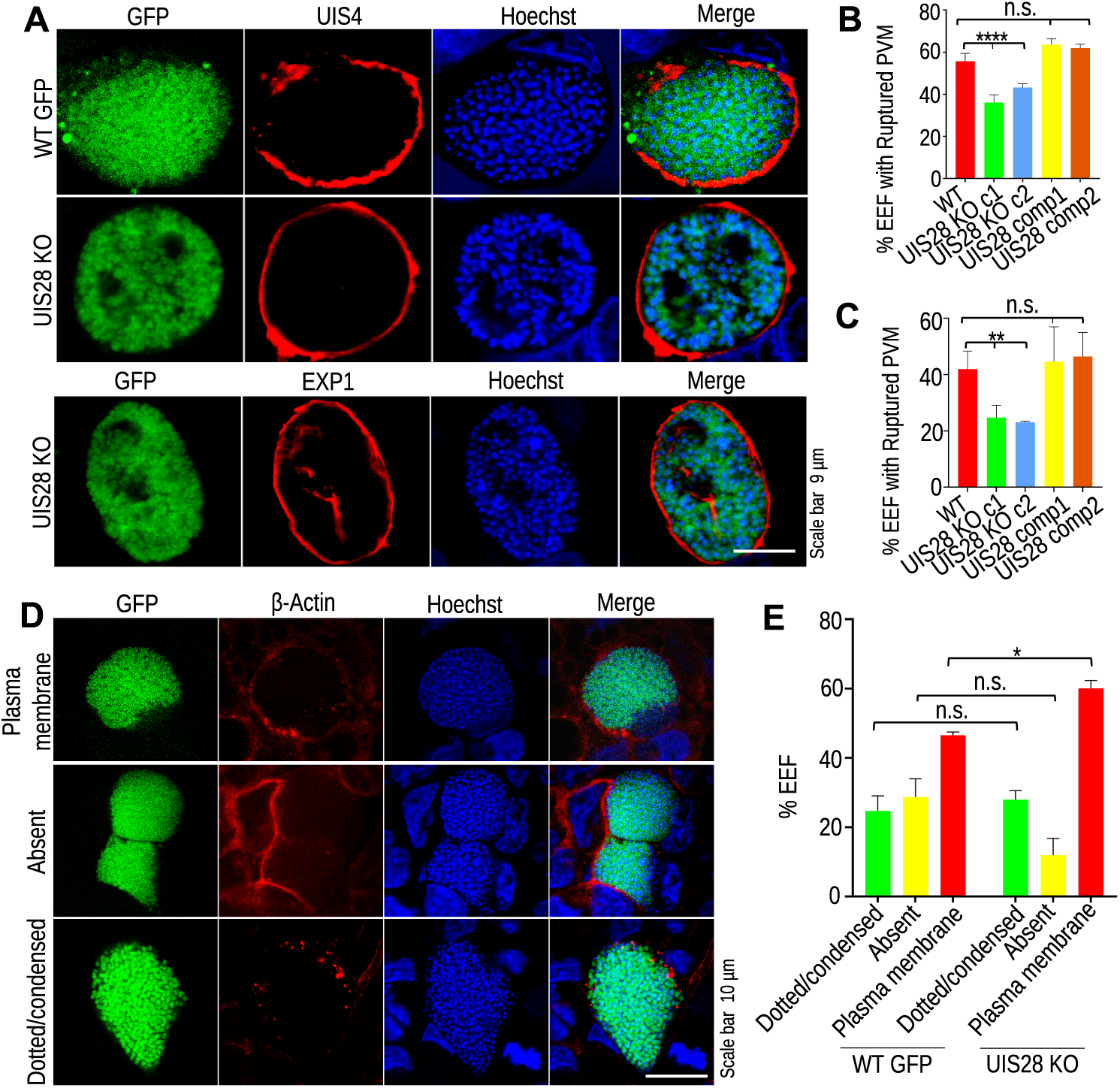
UIS28 KO parasites show a defect in PVM rupture. **(A)** Infected HepG2 cultures fixed at 65 hpi, were immunostained with the PVM markers anti-UIS4 and anti-EXP1 antibodies. Nuclei were stained with Hoechst. (**B and C)** Intact and ruptured PVMs were quantified, and we detected a significantly greater number of EEFs with intact PVMs in UIS28 KO than in WT GFP immunostained with anti-UIS4 (****P<0.0001) and anti-EXP1 (**P=0.0068). The data were obtained from multiple coverslips and are presented as the mean ± SD. Statistical significance was determined by one-way ANOVA. **(D)** Infected cultures fixed at 65 hpi were immunostained with an anti-β-actin antibody. Nuclei were stained with Hoechst. Different types of actin localization patterns (condensed/dotted, absent, or associated with the plasma membrane) in infected cells were quantified. **(E)** Quantification of the actin localization pattern in infected cells. We detected a greater percentage of plasma membrane-associated actin in KO parasites than in WT parasites (*P=0.0162, Students t test). The other actin patterns were similar in both groups. A total of 72 (WT GFP) and 112 (UIS28 KO) EEFs were analyzed. Data were pooled from two independent experiments and are shown as the mean ± SD.

## Discussion

*Plasmodium* parasites escape from hepatocytes by forming structures called merosomes [19]. The process is completed in two subsequent steps. The first phase involves disruption of the PVM, followed by rupture of the host cell plasma membrane. Lipids and phospholipids are abundant in parasite membranes. Parasites substantially modify these membranes during their release from hepatocytes [2]. While proteases [30], kinases [32,33], and phospholipases [20,31] are involved in parasite egress from the hepatocytes, the role of a lipase, which is a prerequisite for lipid breakdown and membrane disruption, remains unknown.

Here, for the first time, we confirmed that a *P. berghei* protein, UIS28, possesses lipase activity. In support of the function of this lipase (UIS28) in PVM disruption, the protein was found to be located at the PVM and detectable at 48–65 hpi, a period prior to and during the PVM rupture. UIS28-deficient parasites are defective in liver stage merozoite egress due to impaired PVM disruption. While studies in Listeria have revealed the importance of two phospholipases (phosphatidylinositol-phospholipase C and phosphatidylcholine-phospholipase C) in membrane disruption [34,35], we describe the exact function of a class-III lipase, UIS28, in *Plasmodium* in the destruction of the PVM. Several bacterial proteases and phospholipases have been found to be involved in the destruction of structural or cytoskeletal proteins that maintain the membrane integrity and thereby help in the exit process [36]. The PVM lacks the underlying cytoskeleton found in the host cell plasma membrane. After PVM rupture, host actin is released into the parasite cytosol [37]. We detected a lower proportion of UIS28 KO parasites whose cytosol exhibited host actin, indicating that UIS28 lipase activity is not only important for PVM rupture but also possibly involved in the destruction of host actin.

UIS28 is expressed in the salivary gland sporozoites. However, sporozoites depleted of UIS28 still exhibit normal hepatocyte invasion, indicating that the expression of the gene in salivary gland sporozoites has a redundant function. This is similar to several UIS proteins, including UIS3, UIS4, and SPELD, which are highly expressed in salivary gland sporozoites but play essential roles during liver stage development [18,38,39]. A member serine repeat antigen (SERA) family, and several phospholipases have been reported to be crucial for the egress of parasites [20,28,30,31,40,41]. Cysteine protease activity is crucial for PVM rupture in *Plasmodium* parasites. The egress of merozoites from hepatocytes was completely blocked by the cysteine protease inhibitor E64, indicating the essential role of these enzymes in egress [19]. The cysteine proteases involved in this process are encoded by genes of a multigene family called SERA [26,42–44]. Kinases play crucial roles in activating the proteases implicated in parasite egress [45]. As kinases and proteases do not have membranolytic activity, they possibly activate downstream effector molecules such as lipases to promote the membrane lytic activity. The members of the SERA family localize to the PVM in hepatocytes, and most of them are expressed during late liver stage development, suggesting their possible role in the initiation of the egress process [26,46]. Therefore, we hypothesize that UIS28 is possibly a downstream target of these kinases and proteases that initiates the PVM rupture. Electron tomography studies of blood stage parasites have suggested that the PVM is not permeabilised via major membrane breakage but rather due to the formation of small localized pores [47]. While *Plasmodium* blood and liver stages show distinct differences in their egress mechanisms, they also share some striking similarities. It is therefore tempting to speculate that UIS28 is possibly involved in PVM rupture by digesting membrane lipids and creating small pores in the PVM.

Disruption of UIS28 did not affect the growth or development of parasites in the blood, mosquito, or liver stages. However, several mature UIS28 KO EEFs presented defective PVM rupture and egress of hepatic merozoites. Similarly, LISP1, SERA4, PbPL and PbPLa1 mutant parasites fail to egress efficiently from infected hepatocytes [20,29–31]. Our study revealed that UIS28 is involved in disrupting the PVM. However, even in UIS28 KO parasites, a proportion of EEFs can still disrupt the PVM, suggesting that other mechanisms are also involved. *Plasmodium* parasites can utilize multiple strategies to exit host cells, as evidenced by the fact that even when proteins involved in egress, such as LISP1 [29], PbPL [20], PbSERA4 [30], and PbPla1 [31] are absent, the parasite still demonstrates some ability to exit the host cells.

In conclusion, our study identified UIS28 as the first *Plasmodium* lipase involved in PVM disruption and egress of hepatic merozoites. Notably, parasites lacking UIS28 do not exhibit a complete egress defect, indicating that additional factors may compensate for the absence of UIS28. This partial phenotype offers a valuable opportunity to dissect the molecular mechanism of the parasite egress and to identify other molecular players that act in combination to facilitate the parasite exit from the host cells.

## Materials and methods

### Ethics statement

All animal work was performed according to the CSIR-CDRI Institutional Animal Ethics Committee (approval no: IAEC/2018/03, IAEC/2023/15 and IAEC/2025/33) set under the Committee for the Purpose of Control and Supervision of Experiments on Animals (CPCSEA) guidelines.

### Experimental organisms, parasites and cell lines

All the mice used in this study were bred and kept in the CSIR-CDRI animal facility. Six-to eight-week-old female Swiss albino and C57BL/6 mice were used for *P. berghei* propagation, transmission, and sporozoite infection, respectively. *P. berghei* ANKA (MRA-311) and *P. berghei* ANKA wild type GFP (MRA-867 507 m6cl1) were obtained from BEI Resources, USA. *Anopheles stephensi* mosquitoes were reared in the insectary facility of CSIR-CDRI. *P. berghei* sporozoites were obtained via infection of female *A. stephensi* mosquitoes as previously described [48]. Human liver hepatocellular carcinoma (HepG2) cells were cultured in DMEM supplemented with 10% FBS, 0.2% NaHCO3, 1% sodium pyruvate, and 1% penicillin‒streptomycin at 37°C with 5% CO2.

### UIS28 amino acid sequence analysis

UIS28 (PBANKA_1017500) amino acid sequence was retrieved from PlasmoDB and analyzed using BLAST (RID-2U5PYNGN016) [49], InterPro [50], SMART [51,52], CD-Search (NCBI) [53–55], TMHMM [56], ScanProsite tool [57]. Protein structure prediction was performed using iTASSER server for protein 3D modelling [58,59].

### Expression and purification of recombinant UIS28

The sequence encoding the catalytic domain of UIS28 was amplified from *P. berghei* cDNA via primers 2155/2156 (primers are detailed in Table S2). The amplified product was cloned in frame with GST into the pGEX-5X-1 vector at the EcoRI restriction site. The correct sequence and expression cassette were confirmed via DNA sequencing. The expression of the GST-UIS28 fusion protein was induced via 0.2 mM IPTG for 16 h at 20°C in Rosetta (DE3) cells as previously described [60]. After induction, the cells were harvested and lysed in a buffer containing 10 mg/mL lysozyme and protease inhibitors. The lysate was sonicated at 20Hz for 5 minutes with 30 seconds pulse-on and 20 seconds pulse-off, followed by incubation with 0.1% Triton-X-100 on ice for 30 min and subsequent centrifugation at 12,000 × g. The supernatant was passed through glutathione Sepharose beads, and the bound protein was eluted with glutathione. The purified protein was desalted via a 10 kDa MWCO centrifugal concentrator (Amicon).

### In vitro lipase assay

The lipase assay was performed using a lipase assay kit (MAK046, Sigma-Aldrich) following the manufacturer’s instructions. IPTG-induced secondary culture containing 2 × 10^6^ bacterial cells was homogenized in 4 volumes of ice-cold lipase assay buffer. The assay is based on a series of coupled enzymatic reactions in which glycerol is used as a standard. The amount of endogenous glycerol in the sample was measured, and the net amount of glycerol generated by lipase was calculated by subtracting it from the blank. The amount of glycerol produced is proportional to the enzymatic activity of the lipase. The lipase activity was expressed as milliunits/mL, where 1 unit of lipase generates 1 μmole of glycerol per minute at 37°C.

### Generation of UIS28 m-Cherry-tagged, KO and complement parasite lines in *P. berghei*

To endogenously tag the UIS28 gene with 3XHA-mCherry a DCO strategy was used as previously described [61]. Two fragments, F1 (0.55 kb) and F2 (0.57 kb), encompassing the C-terminus of the gene and 3’UTR, were amplified via primers 2052/2053 and 2050/2051 and cloned at *the* XhoI/BglII and NotI/AscI sites, respectively. The construct was linearized via XhoI/AscI and transfected into *P. berghei* ANKA achizonts as previously described [62]. To disrupt the UIS28 (PBANKA_1017500) target gene locus, we employed a double-crossover (DCO) homologous recombination. Two homologous fragments, F3 (0.66 kb) and F4 (0.54 kb), from the flanking regions of the target gene were amplified via the primer pairs 1758/1759 and 1760/1761, respectively. The amplified fragments F3 and F4 were then sequentially cloned into the pBC-GFP-hDHFR:yFCU vector at the *Xho*I/*Sal*I and *Not*I/*Asc*I sites, respectively. The resulting construct was linearized via *Xho*I/*Asc*I restriction digestion before being transfected into *P. bergehi* ANKA schizonts. The UIS28-complemented parasite line was subsequently generated by amplifying fragment F5 (4.3 kb), which encompasses the PbUIS28 ORF and the 5’ and 3’ UTRs. The fragment was then transfected into UIS28 KO schizonts. The UIS28^comp^ parasites were negatively selected via the 5-fluorocytosine (5-FC) drug [63], whereas the UIS28 KO and UIS28-3XHA mCherry parasites were selected via the pyrimethamine drug (Sigma-Aldrich, 46706). Parasite clonal lines were generated by limiting dilution. All the genetic modifications of the parasites were confirmed via diagnostic PCR using the primer pairs 1834/1225 and 1215/1835 for UIS28 KO parasites, and 2154/1218 and 1215/1835 for UIS28^mch^ parasites. The absence of the UIS28 ORF in UIS28 KO and ORF restoration in UIS28^comp^ were confirmed via the primer set 1836/1837. All the oligonucleotide sequences are listed in Table S2.

### Western blot analysis

UIS28-3XHA-mCherry sporozoites and induced GST-UIS28 bacterial cultures were lysed in Laemmli sample buffer (Bio-Rad, 1610747), resolved via 7.5% SDS-PAGE and transferred onto a nitrocellulose membrane (Bio-Rad, USA) by electroblotting as previously described [64]. The membrane was then probed with mouse anti-mCherry (dilution 1:1,000, 1C51,ThermoFisher), mouse anti-CSP (1:1,000) [65] or goat anti-GST(dilution 1:5,000, 27457701V, Sigma-Aldrich) antibodies, followed by incubation with horseradish-peroxidase (HRP)-conjugated anti-rabbit or anti-mouse (dilution 1:5,000, Amersham Biosciences, United Kingdom, NA934V/NA931V) or anti-goat (dilution 1:10,000,Thermoscientific, A15999) secondary antibodies. The protein bands were then visualized via ECL Chemiluminescent Substrate (Bio-Rad, 170-5060), and images were captured via the ChemiDoc XRS+ System (Bio-Rad, USA).

### Analysis of UIS28 KO parasite development in asexual blood and mosquito stages

To analyze the asexual blood stage propagation, an equal number of infected red blood cells (iRBCs) of the WT GFP, UIS28 KO, and UIS28^comp^ parasite lines were intravenously (i.v.) injected into Swiss albino mice (5 mice/group). Parasitemia was monitored for a period of five days via Giemsa-stained blood smear. The infection of female *A. stephensi* was initiated by allowing mosquitoes to probe for the blood meal in infected Swiss albino mice. The infected mosquito cages were kept in an environmental chamber maintained at 19°C with 80% relative humidity. To analyze the ookinete development, infected mosquito midguts were dissected 24 h post-blood meal, and the ookinetes were observed under a fluorescence microscope as previously described [66]. The number of ookinetes was quantified via a hemocytometer. On days 14 and 18 post-blood meal, mosquito midguts and salivary glands were dissected and homogenized, and sporozoite numbers were enumerated via a hemocytometer. Additionally, batches of infected midguts and salivary glands were visualized under a fluorescence microscope to evaluate the number of oocysts and sporozoite load in the oocysts and salivary glands.

### In vivo infection of sporozoites in mice

To evaluate the in vivo infection, 5,000 sporozoites of WT GFP, UIS28 KO, and UIS28^comp^ parasites were intravenously injected into different groups of C57BL/6 mice. The progression of parasites to initiate the blood stage infection was monitored daily via microscopic examination of Giemsa-stained blood smear. To determine the parasite burden in the liver, two groups of C57BL/6 mice were inoculated with 5,000 WT GFP or UIS28 KO sporozoites, and the livers were harvested and homogenized in 10 mL of RNAiso Plus reagent (Takara, 9108). Total RNA was isolated from the liver homogenate by following the manufacturer’s instructions. The RNA was reverse transcribed to synthesize a complementary DNA (cDNA) strand as previously described [67]. Pb18S rRNA was amplified from cDNA via the primers 1195/1196 and quantified via real-time PCR using SYBR Premix (Takara Bio, RR420A) in a real-time PCR machine (CFX Opus 96 real-time PCR system; Bio-Rad). The transcripts were normalized to the amount of mouse GAPDH via amplification using primers 1193/1194. The primers used in the study are listed in Table S2.

### In vitro analysis of liver stage development

Human hepatocellular carcinoma (HepG2) cells were cultured in DMEM supplemented with 10% FBS at 37°C and 5% CO2. One day prior to sporozoite infection, 5.5 x 10^4^ cells/well were seeded onto collagen-coated glass coverslips in a 48-well cell culture plate. For detached cell analysis, 1 × 10^5^ cells/well were seeded into a collagen-coated 24-well cell culture plate. Salivary gland sporozoites (5 x 10^3^/well in 48-well plates and 3.0 x 10^4^/well in 24-well plates) were added to the cultures and maintained as previously described [68]. Cultures in 48-well plates were terminated at 45 and 65 hpi and fixed with 4% paraformaldehyde (PFA), as previously described. Cultures in a 24-well plates were allowed to grow until 65-78 hpi and visualized under a light microscope to assess the detached cell formation. For detached cell quantification, the culture supernatant was collected, and the number of detached cells was counted via a hemocytometer. The ability of detached cells to infect RBCs was monitored by injecting them intravenously into Swiss albino mice. The infection of the mice was monitored via Giemsa-stained blood smear.

### Sporozoite invasion assay

To analyze the invasion efficiency of salivary gland sporozoites, HepG2 cells were infected as described above. The cultures were harvested at 1.5 h post-sporozoite addition and fixed with PFA. To quantify the sporozoites inside vs. outside, a differential immunofluorescence assay was performed using an anti-CSP antibody [65] as previously described [69].

### Generation of anti-EXP1 antibody

To visualize PVM, the EXP1 antibody was generated in the rabbit. KLH-conjugated peptides of EXP1 (KNKHGKTGSKNVIKKP-Cys) was synthesized by GL Biochem (Shanghai) Ltd. The peptide was used for immunization in complete Freund’s adjuvant for priming and in incomplete Freund’s adjuvant for the booster. The rabbit was bled, and the serum was collected for further analysis.

### Immunofluorescence assays

Salivary gland sporozoites were allowed to settle onto 12-well slides (Thermo Fisher Scientific, USA), air-dried, and fixed with 4% PFA (Sigma–Aldrich, HT5012) as previously described [48]. Sporozoites were permeabilized with 0.1% Triton-X-100 (Sigma–Aldrich, T8787) for 10 minutes at room temperature. Fixed infected HepG2 cultures were permeabilized with chilled methanol for 20 minutes at 4°C. Non-specific binding sites were blocked by incubation with 1% BSA/PBS for 1 h. Sporozoites were incubated with mouse anti-mCherry (dilution 1:500, 1C51,ThermoFisher), and rabbit anti-TRAP [70] antibodies. HepG2 cultures were incubated with mouse anti-mCherry (dilution 1:500, 1C51,ThermoFisher), rabbit anti-UIS4 [16], mouse anti-MSP1 (mouse monoclonal; diluted 1:5,000) [71], and rabbit anti-EXP1 (polyclonal; diluted 1:1,000), mouse anti-CSP [65] (diluted 1:1,000) and mouse anti-β actin (Invitrogen-MA1-140, diluted 1:1000). The signals of the primary antibodies were revealed via Alexa Fluor 488-conjugated or Alexa Fluor 594-conjugated secondary antibodies (Invitrogen, diluted 1:1,000). Hoechst 33342 (Sigma–Aldrich, 41399) was used to stain and visualize the DNA in all IFA experiments. Slides were mounted using Prolong Diamond antifade reagent (Invitrogen, P36970). Representative EEF images were acquired via FV1000 software on a confocal laser scanning microscope (Olympus BX61WI) via a UPlanSAPO 100x (NA 1.4, oil) or 63× (NA 0.25, oil).

### Statistical analysis

All the statistical analyses were carried out via GraphPad Prism software (version 9.0). The data are reported either as the mean ± SEM or mean ± SD, as indicated. Differences between the two experimental groups were assessed via an unpaired two-tailed Student’s t test. For comparisons between more than two experimental groups, one-way ANOVA was used. A p value of <0.05 was considered to indicate statistical significance.

## Acknowledgments

We thank BEI Resources, USA, for the parasite strains. We thank Dr. Photini Sinnis (Johns Hopkins University, USA) and Dr. Anthony A. Holder (The Francis Crick Institute, UK) for their anti-UIS4 and anti-MSP1 antibodies, respectively. We acknowledge the CSIR-CDRI’s Intravital microscopy facility. The University Grants Commission and Council of Scientific and Industrial Research, Government of India, research fellowships supported RD and PM. This study was supported by CSIR-CDRI in-house funding. This manuscript is CDRI communication no. 91/2025/SM.

## Data availability

The data supporting the findings of this study are available within the paper and its supplementary information files. All the datasets and raw data analyzed during the current study will be made available upon request.

## Author Contributions

Conception and design of study: RD, SM; Acquisition of data: RD, SM; Analysis of data: RD, SM; Methodology: RD, PM, SA; Drafting and revisions of the Manuscript: RD, SM. All authors have read and approved the final version of the manuscript.

Declaration of Interests

The authors declare that they have no competing interests.

## Supplementary Material

**Fig. S1.**
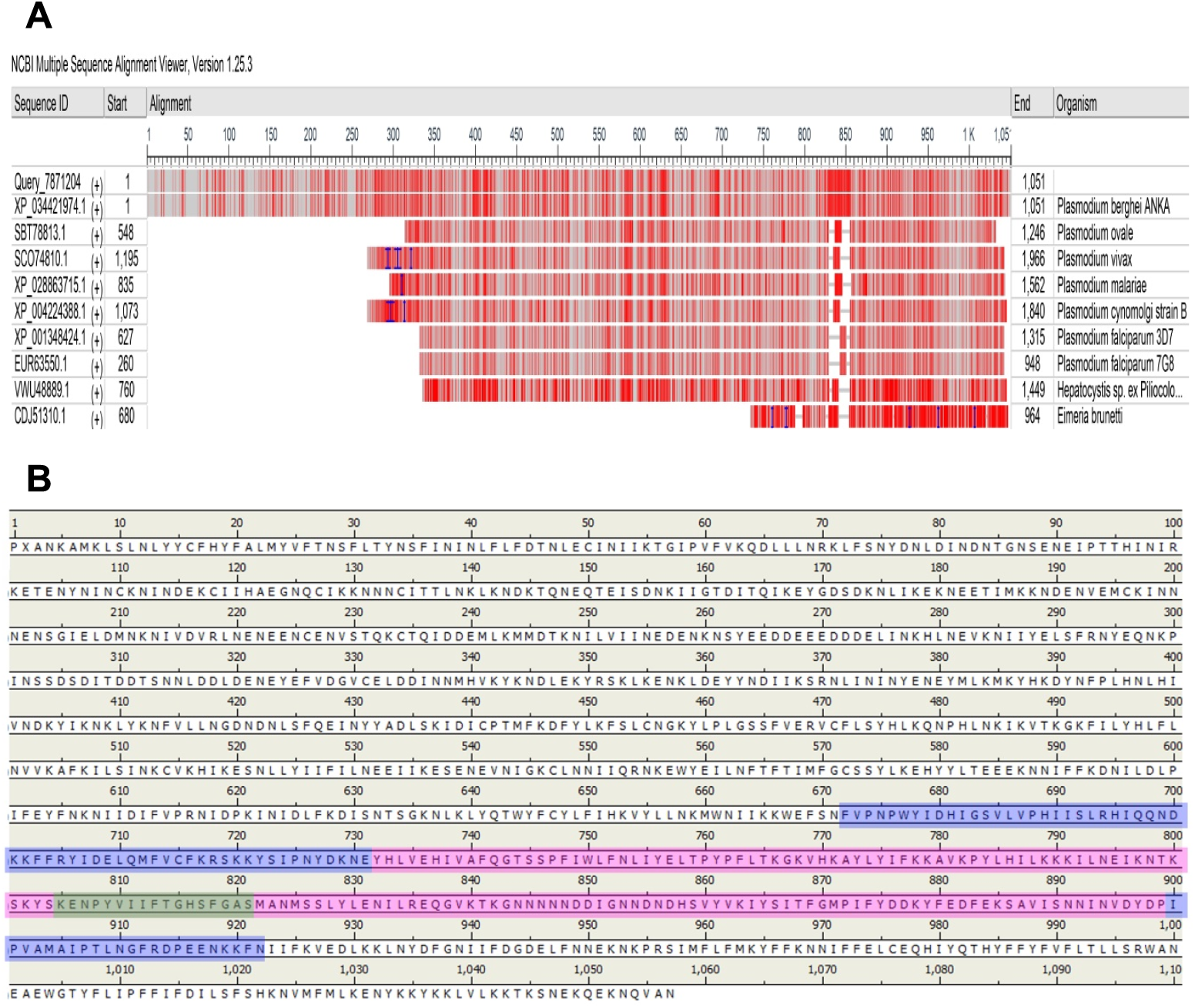
(A) NCBI-BLAST multiple sequence alignment of UIS28 showing the sequence similarity. **(B)** The primary structure of UIS28 highlighting the long α/β-hydrolase fold spanning from 672-923 (blue) with intervening class 3 lipase domain (732-899, pink). The catalytic stretch with key the residues ‘PYVIIFTGHSFGAS’ in the lipase domain (green) holds the catalytic serine-811.

**Fig. S2.**
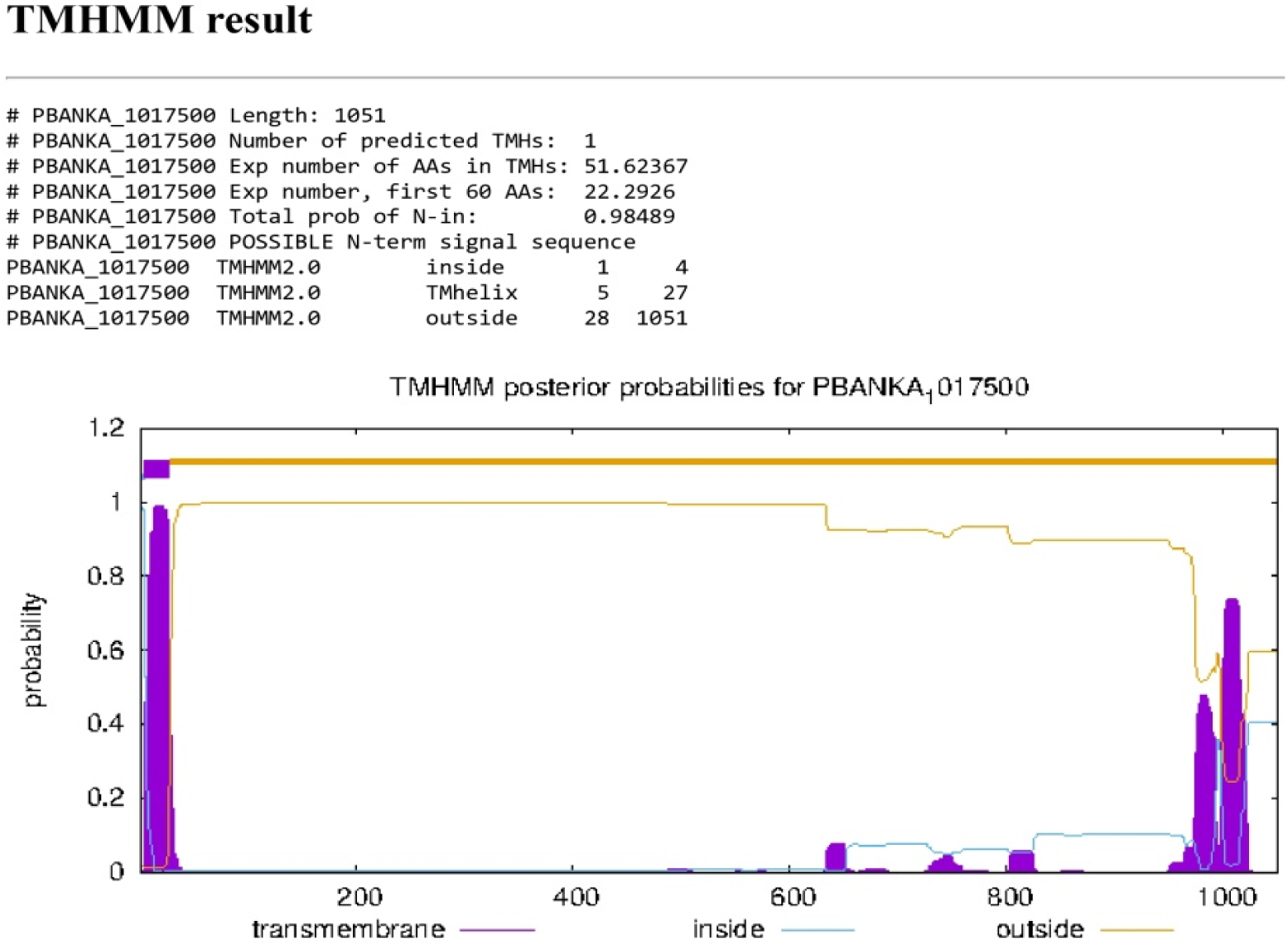
The TMHMM result shows the multiple transmembrane helices (violet), followed by a long stretch of outer membrane regions (orange line) and intervening short cytoplasmic regions (blue line) of UIS28 domains.

**Fig. S3.**
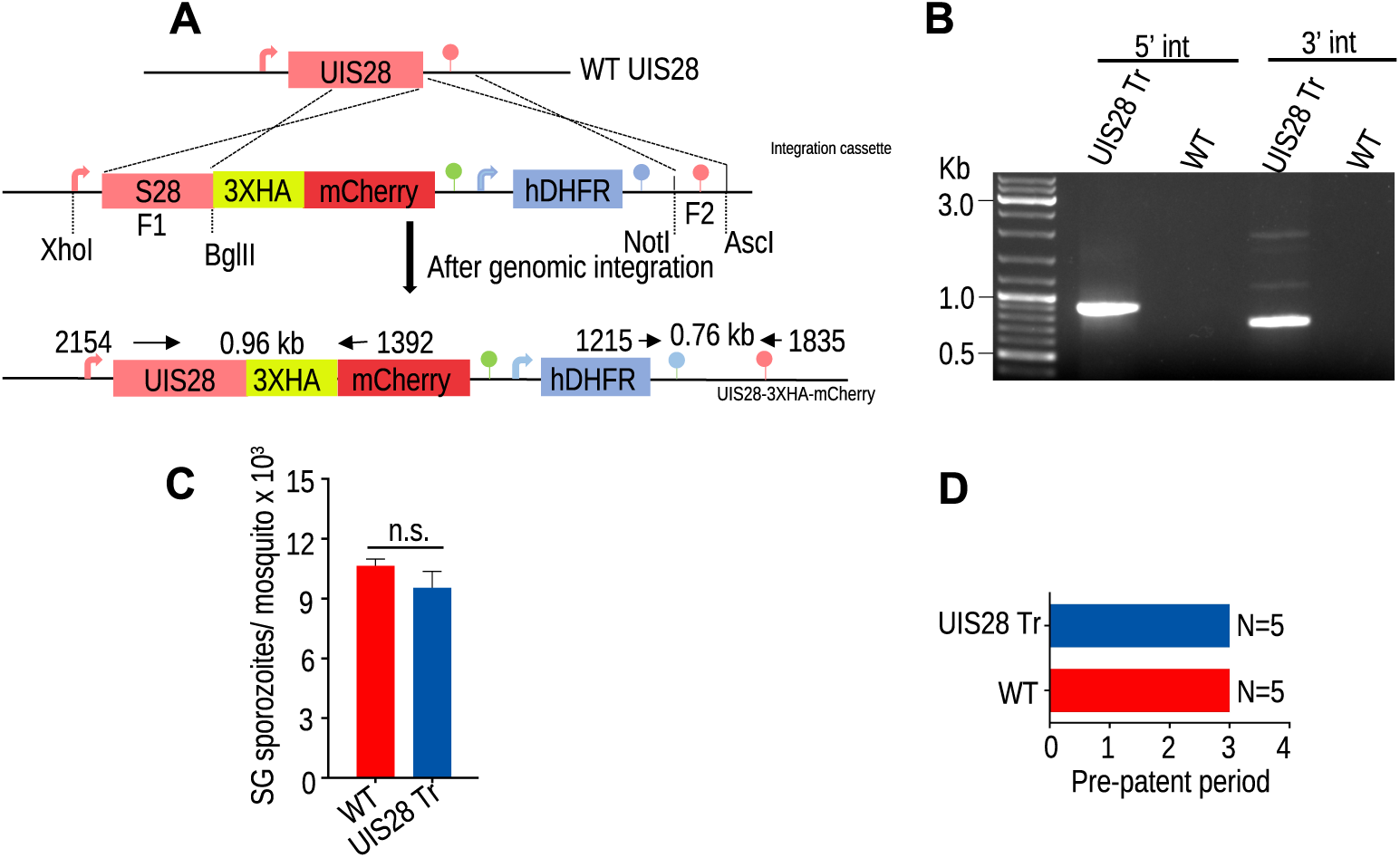
Generation of UIS28-3XHAmCherry transgenic parasites. **(A)** Schematic representation of the gene tagging strategy with 3XHAmCherry. **(B)** Diagnostic PCR with primer sets 2154/1392 and 1215/1835 confirmed the correct 5’ and 3’ integrations, respectively. **(C)** Comparable numbers of salivary gland sporozoites were produced in WT and UIS28-3XHAmCherry transgenic parasites (P=0.2223, Student’s t test). The data are shown as mean ± SD. **(D)** UIS28-3XHA-mCherry sporozoites infect C57BL/6 mice normally. N, number of mice.

**Fig. S4.**
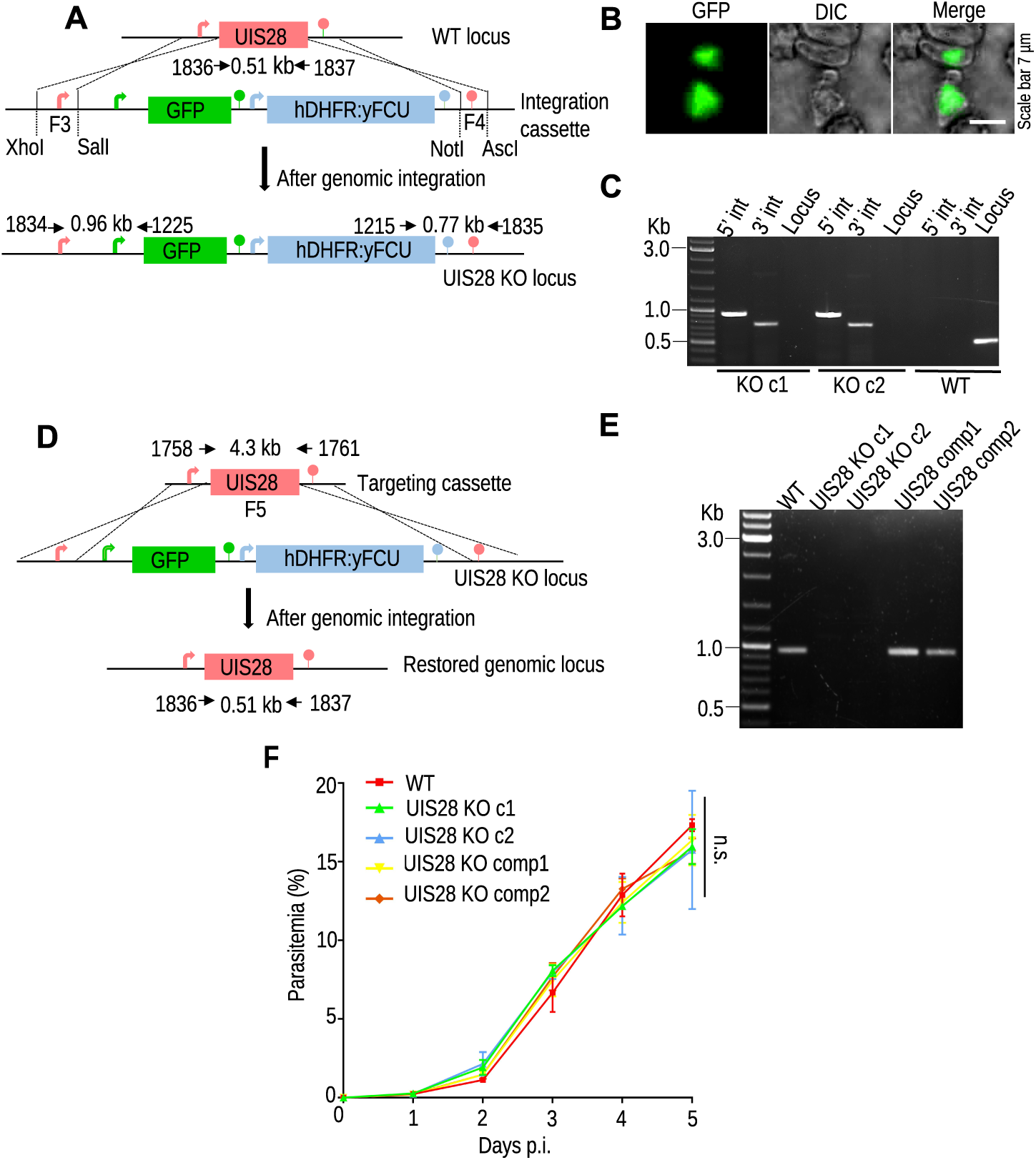
UIS28 is not required for asexual blood stage propagation. **(A)** Schematic showing the gene deletion strategy used for UIS28. After double-crossing over the homologous recombination, the WT locus was replaced by an integration cassette. **(B)** GFP-expressing UIS28 KO blood stage parasites. **(C)** Confirmation of correct genomic integration of the targeting cassette via site-specific integration PCR using primer sets 1834/1225 and 1215/1835 for 5’ and 3’ integration, respectively. WT gDNA was used as a negative control. The primer set 1836/1837 was used to confirm the absence of the WT UIS28 locus in the KO parasites. **(D)** Complementation strategy to reintroduce the WT UIS28 gene at the KO locus. **(E)** Restoration of the WT UIS28 locus was confirmed via diagnostic PCR using primer pair 1836/1837. **(F)** Propagation of the asexual blood stage in the WT, UIS28 KO clonal lines, and UIS28 complement parasite lines (5 mice/group). No significant difference in asexual blood propagation was observed (P=>0.9999, one-way ANOVA). Data from two independent experiments are shown as the mean ± SEM.

**Fig. S5.**
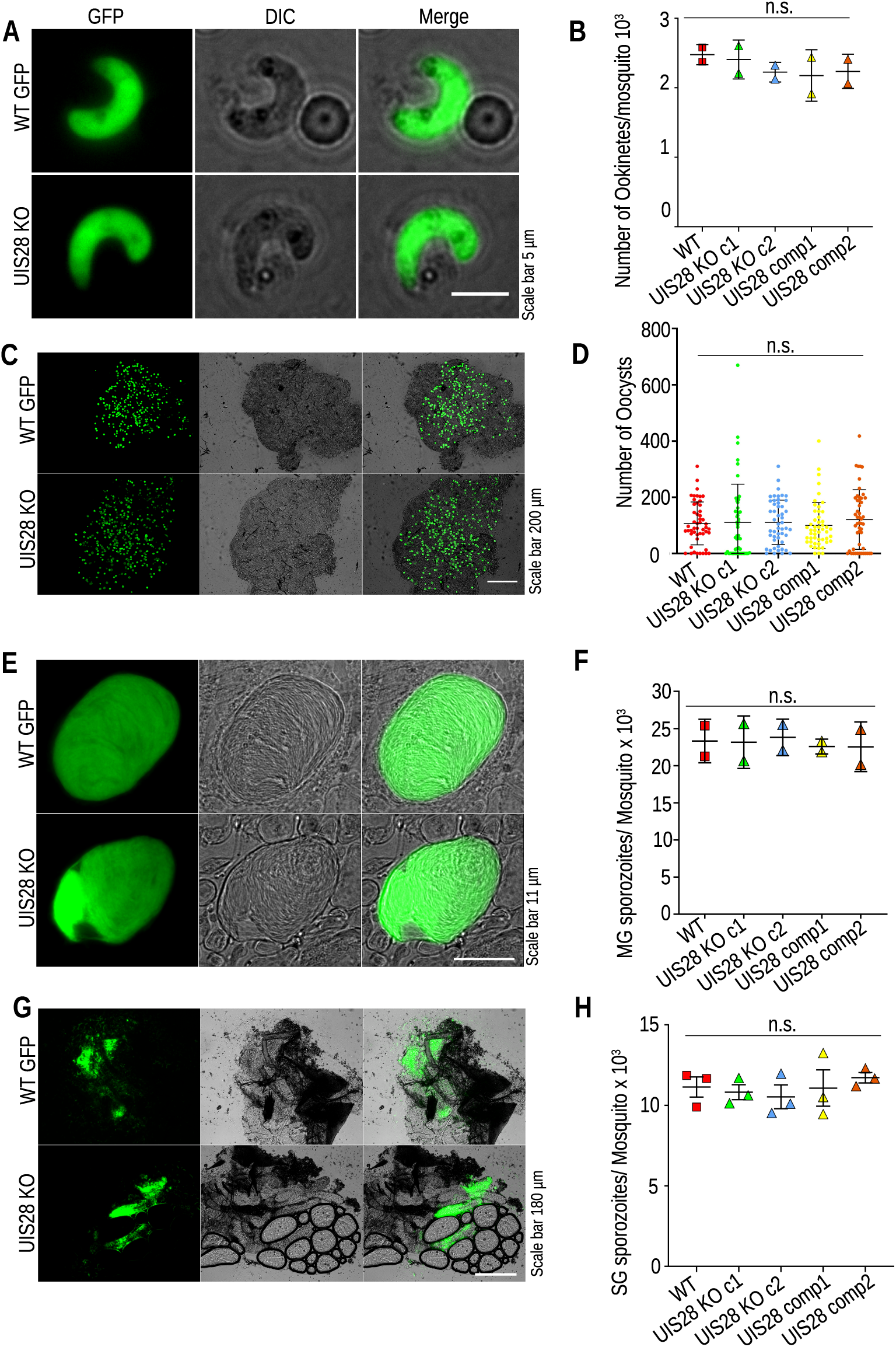
UIS28 KO parasites develop normally during the mosquito stages. **(A)**Representative images showing normal ookinete morphology in UIS28 KO and WT parasites. **(B)** Comparable ookinete numbers in WT GFP and UIS28 KO parasites (P=0.7181). Data from 76 (WT GFP), 61(UIS28 KO c1), 41 (UIS28 KO c2), 38 (UIS28 comp1) and 50 (UIS28 comp2) mosquitoes are shown. **(C)** Similar patterns of midgut oocyst development in WT GFP and UIS28 KO infected mosquitoes. **(D)** The number of oocysts produced in UIS28 KO-infected mosquito midguts was comparable to that produced in WT mosquito midguts (P=0.8780). **(E)** Sporogony pattern in WT GFP and UIS28 KO oocysts. **(F)** Quantification of the number of midgut sporozoites. No difference was observed between control and KO parasites (P=0.9874). A total of 42 (WT GFP), 85 (UIS28 KO c1), 46 (UIS28 KO c2), 40 (UIS28 comp1), and 48 (UIS28 comp2) mosquito midguts were analyzed **(G).** WT GFP and UIS28 KO-infected mosquito salivary glands presented similar sporozoite loads. **(H)** Quantification of salivary gland sporozoites revealed a comparable number of sporozoites in the control and KO groups (P=0.8141). Data from 190 (WT GFP), 198 (UIS28 KO c1), 296 (UIS28 KO c2), 244 (UIS28 comp1), and 207 (UIS28 comp2) mosquitoes. The data are presented as mean ± SEM and were analyzed via one-way ANOVA.

**Fig. S6.**
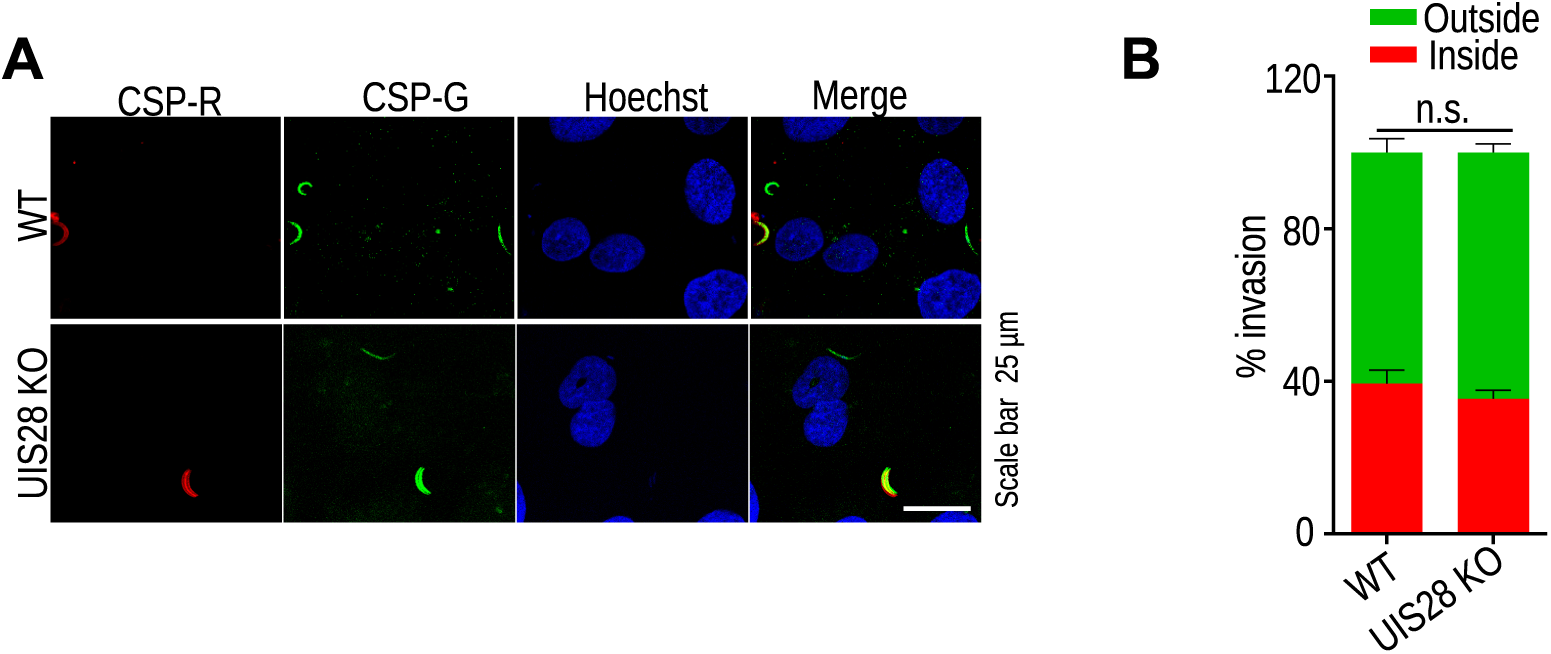
UIS28 KO sporozoites efficiently invade hepatocytes. **(A)** HepG2 cultures infected with WT GFP and UIS28 KO sporozoites were fixed at 1.5 hpi. The coverslips were immunostained with anti-CSP antibody before and after permeabilization. The sporozoites that failed to invade hepatocytes were stained red, while green signifies the total number of sporozoites. **(B)** The percent invasion of WT GFP and UIS28 KO sporozoites was not significantly different between WT and KO parasites (P=0.0982, Student’s t test). The data are expressed as the mean ± SD and were pooled from two independent experiments.

**Table S1.** Sequence alignment revealed that the UIS28 protein is conserved in malaria parasites. The numbers are similarity percentage values.

| Organism | Accession ID | Query Coverage | % Identity |
| --- | --- | --- | --- |
| *Plasmodium berghei* | XP_034421974.1 | 100 | 100 |
| *Plasmodium ovale* | SBT78813.1 | 64.117 | 79.97 |
| *Plasmodium vivax* | SCO74810.1 | 57.449 | 75.63 |
| *Plasmodium malariae* | XP_028863715.1 | 61.015 | 78.64 |
| *Plasmodium cynomolgi* | XP_004224388.1 | 57.614 | 75.63 |
| *Plasmodium falciparum* 3D7 | XP_001348424.1 | 53.305 | 74.4 |
| *Plasmodium falciparum* K1 | EUR63550.1 | 53.305 | 74.4 |
| Hepatocystis sp. Ex Piliocolobus tephrosceles, Lipase putattive | VWU48889.1 | 67 | 50.7 |
| Eimeria brunette hypothetical protein | CDJ51310.1 | 29 | 26.3 |
| Bacillus sp. Lipase family |  | 8 | 33 |
| Human |  |  | No similarity |

**Table S2.**
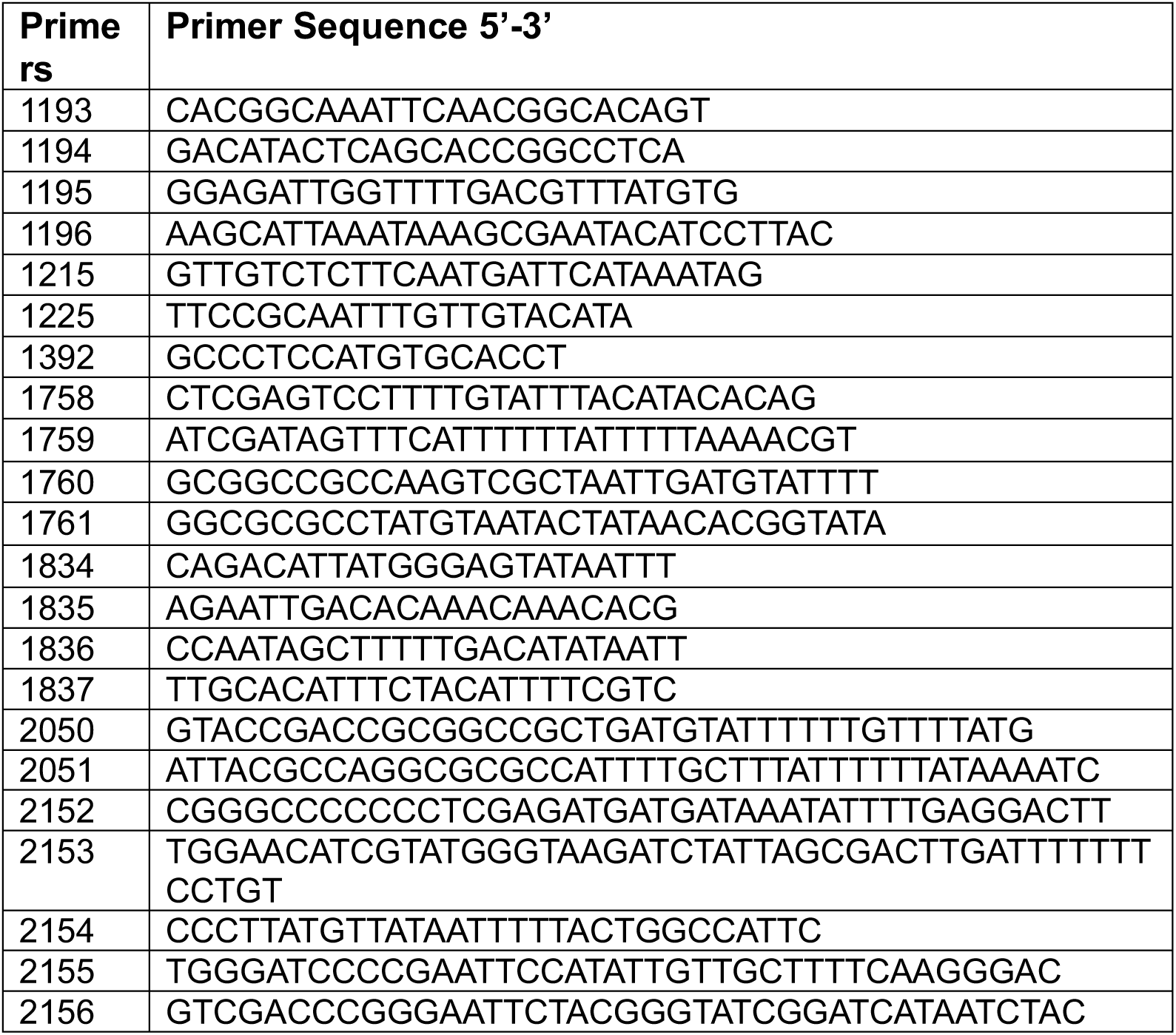
Primers used in this study.

